# Expanding the stdpopsim species catalog, and lessons learned for realistic genome simulations

**DOI:** 10.1101/2022.10.29.514266

**Authors:** M. Elise Lauterbur, Maria Izabel A. Cavassim, Ariella L. Gladstein, Graham Gower, Nathaniel S. Pope, Georgia Tsambos, Jeff Adrion, Saurabh Belsare, Arjun Biddanda, Victoria Caudill, Jean Cury, Ignacio Echevarria, Benjamin C. Haller, Ahmed R. Hasan, Xin Huang, Leonardo Nicola Martin Iasi, Ekaterina Noskova, Jana Obšteter, Vitor Antonio Corrêa Pavinato, Alice Pearson, David Peede, Manolo F. Perez, Murillo F. Rodrigues, Chris C. R. Smith, Jeffrey P. Spence, Anastasia Teterina, Silas Tittes, Per Unneberg, Juan Manuel Vazquez, Ryan K. Waples, Anthony Wilder Wohns, Yan Wong, Franz Baumdicker, Reed A. Cartwright, Gregor Gorjanc, Ryan N. Gutenkunst, Jerome Kelleher, Andrew D. Kern, Aaron P. Ragsdale, Peter L. Ralph, Daniel R. Schrider, Ilan Gronau

**Affiliations:** Department of Ecology and Evolutionary Biology, University of Arizona, Tucson AZ 85719, USA; Department of Ecology and Evolutionary Biology, University of California, Los Angeles, Los Angeles CA, USA; Embark Veterinary, Inc., Boston MA 02111, USA; Section for Molecular Ecology and Evolution, Globe Institute, University of Copenhagen, Denmark; Institute of Ecology and Evolution, University of Oregon, Eugene OR 97402, USA; School of Mathematics and Statistics, University of Melbourne, Australia; AncestryDNA, San Francisco CA 94107, USA; 54Gene, Inc., Washington DC 20005, USA; Université Paris-Saclay, CNRS, INRIA, Laboratoire Interdisciplinaire des Sciences du Numérique, UMR 9015 Orsay, France; School of Life Sciences, University of Glasgow, Glasgow, UK; Department of Computational Biology, Cornell University, Ithaca NY, USA; Department of Cell and Systems Biology, University of Toronto, Toronto ON, Canada; Department of Biology, University of Toronto Mississauga, Mississauga ON, Canada; Department of Evolutionary Anthropology, University of Vienna, Vienna, Austria; Human Evolution and Archaeological Sciences (HEAS), University of Vienna, Vienna, Austria; Department of Evolutionary Genetics, Max Planck Institute for Evolutionary Anthropology, Leipzig, Germany; Computer Technologies Laboratory, ITMO University, St Petersburg, Russia; Agricultural Institute of Slovenia, Department of Animal Science, Ljubljana, Slovenia; Entomology Department, The Ohio State University, Wooster OH, USA; Department of Genetics, University of Cambridge, Cambridge, UK; Department of Zoology, University of Cambridge, Cambridge, UK; Department of Ecology, Evolution, and Organismal Biology, Brown University, Providence RI, USA; Center for Computational Molecular Biology, Brown University, Providence RI, USA; Department of Genetics and Evolution, Federal University of Sao Carlos, Sao Carlos 13565905, Brazil; Department of Genetics, Stanford University School of Medicine, Stanford CA 94305, USA; Department of Cell and Molecular Biology, National Bioinformatics Infrastructure Sweden, Science for Life Laboratory, Uppsala University, Husargatan 3, SE-752 37 Uppsala, Sweden; Department of Integrative Biology, University of California, Berkeley, Berkeley CA, USA; Department of Biostatistics, University of Washington, Seattle WA, USA; Broad Institute of MIT and Harvard, Cambridge MA 02142, USA; Big Data Institute, Li Ka Shing Centre for Health Information and Discovery, University of Oxford, Oxford OX3 7LF, UK; Cluster of Excellence - Controlling Microbes to Fight Infections, Eberhard Karls Universität Tübingen, Tübingen, Baden-Württemberg, Germany; School of Life Sciences and The Biodesign Institute, Arizona State University, Tempe AZ, USA; The Roslin Institute and Royal (Dick) School of Veterinary Studies, University of Edinburgh, Edinburgh EH25 9RG, UK; Department of Molecular and Cellular Biology, University of Arizona, Tucson AZ 85721, USA; Department of Integrative Biology, University of Wisconsin-Madison, Madison WI, USA; Department of Mathematics, University of Oregon, Eugene OR 97402, USA; Department of Genetics, University of North Carolina at Chapel Hill, Chapel Hill NC 27599, USA; Efi Arazi School of Computer Science, Reichman University, Herzliya, Israel

**Author notes:** These authors contributed equally to the paper (Maria Izabel A. Cavassim, Ariella L. Gladstein, Graham Gower, Nathaniel S. Pope, Georgia Tsambos).

## Abstract

Simulation is a key tool in population genetics for both methods development and empirical research, but producing simulations that recapitulate the main features of genomic data sets remains a major obstacle. Today, more realistic simulations are possible thanks to large increases in the quantity and quality of available genetic data, and to the sophistication of inference and simulation software. However, implementing these simulations still requires substantial time and specialized knowledge. These challenges are especially pronounced for simulating genomes for species that are not well-studied, since it is not always clear what information is required to produce simulations with a level of realism sufficient to confidently answer a given question. The community-developed framework `stdpopsim` seeks to lower this barrier by facilitating the simulation of complex population genetic models using up-to-date information. The initial version of `stdpopsim` focused on establishing this framework using six well-characterized model species (Adrion et al., 2020). Here, we report on major improvements made in the new release of `stdpopsim` (version 0.2), which includes a significant expansion of the species catalog and substantial additions to simulation capabilities. Features added to improve the realism of the simulated genomes include non-crossover recombination and provision of species-specific genomic annotations. Through community-driven efforts, we expanded the number of species in the catalog more than three-fold and broadened coverage across the tree of life. During the process of expanding the catalog, we have identified common sticking points and developed best practices for setting up genome-scale simulations. We describe the input data required for generating a realistic simulation, suggest good practices for obtaining the relevant information from the literature, and discuss common pitfalls and major considerations. These improvements to `stdpopsim` aim to further promote the use of realistic whole-genome population genetic simulations, especially in non-model organisms, making them available, transparent, and accessible to everyone.

## Introduction

Population genetics allows us to answer questions across scales from deep evolutionary time to ongoing ecological dynamics, and dramatic reductions in sequencing costs enable the generation of unprecedented amounts of genomic data that can be used to address these questions (Ellegren, 2014). Ongoing efforts to systematically sequence life on Earth by initiatives such as the Earth Biogenome (Lewin et al., 2022) and its affiliated project networks, such as Vertebrate Genomes (Rhie et al., 2021), 10,000 Plants (Cheng et al., 2018) and others (Darwin Tree of Life Project Consortium, 2022), are providing the backbone for enormous increases in the amount of population-level genomic data available for model and non-model species. These data are being used, among other things, in inference of population history and demographic parameters (Beichman et al., 2018), studying adaptive introgression (Gower et al., 2021), providing null expectations for selection scans (e.g. Hsieh et al., 2021), and understanding the implications of deleterious variation in populations of conservation concern (e.g. Robinson et al., 2023). While many of the methods that address these questions were initially developed for a few key model systems such as humans and *Drosophila*, more recent efforts are generalizing these methods to include important factors not initially accounted for, such as inbreeding or selfing (Blischak et al., 2020), skewed offspring distributions (Montano, 2016), and intense artificial selection even for non-model organisms (MacLeod et al., 2013, 2014).

Simulations can be useful at all stages of this work—for planning studies, analyzing data, testing inference methods, and validating findings from empirical and theoretical research. For instance, simulations provide training data for inference methods based on machine learning (Schrider and Kern, 2018) and Approximate Bayesian Computation (Csilléry et al., 2010). They can also serve as baselines for further analyses: for example, simulations incorporating demographic history serve as null models when detecting selection (Hsieh et al., 2016) or seed downstream breeding program simulations (Gaynor et al., 2020). More recently, population genomic simulations have been used to help guide conservation decisions for threatened species (Teixeira and Huber, 2021; Kyriazis et al., 2022).

Increasing amounts of data and sophistication of inference methods have enabled researchers to ask ever more specific and precise questions. Consequently, simulations must incorporate more and more elements of biological realism. Important elements include genomic features such as mutation and recombination rates that strongly affect genetic variation and haplotype structure (Nachman, 2002). The inclusion of these genomic features is particularly important when linked selection is acting upon the patterns of genomic diversity being studied (Cutter and Payseur, 2013). Furthermore, the demographic history of a species—encompassing population sizes and distributions, divergences, and gene flow—can dramatically affect patterns of genomic variation (Teshima et al., 2006). Thus species-specific estimates of these and other ecological and evolutionary parameters (such as those governing the process of natural selection) are important when generating realistic simulations. This presents challenges, especially to new researchers, as it takes a great deal of specialized knowledge not only to code the simulations themselves but also to find and choose appropriate estimates of the parameters underlying the simulation model.

The recently developed community resource `stdpopsim` provides easy access to detailed population genomic simulations (Adrion et al., 2020). It lowers the technical barriers to performing these simulations and reduces the possibility of erroneous implementation of simulations for species with published demographic models. The initial release of `stdpopsim` was restricted to only six well-characterized model species, such as *Drosophila melanogaster* and *Homo sapiens*, but feedback we received from the community identified a widespread desire to simulate a broader range of non-model species, and ideally to incorporate these into the `stdpopsim` catalog for future use. This feedback, and subsequent efforts to expand the catalog, also uncovered a vital need to better understand when it is practical to create a realistic simulation of a species of interest, and indeed what “realistic” means in this context.

This paper reports on the updates made in the current release of `stdpopsim` (version 0.2), and is also intended as a resource for any researcher who wishes to develop chromosome-scale simulations for their own species of interest. We start by describing the central idea behind the standardized simulation framework of `stdpopsim`, and then outline the main updates made to the `stdpopsim` catalog and simulation framework in the past two years. We then provide guidelines for generating population genomic simulations, either for the purpose of using them in one specific study, or with the intent of making the simulations available for future work by adding the appropriate models to `stdpopsim`. Among other considerations, we discuss when a chromosome-scale simulation is more useful than simulations based on either individual loci or generic loci. We specify the required input data, mention common pitfalls in choosing appropriate parameters, and suggest courses of action for species that are missing estimates of some necessary inputs. We conclude with examples from two species recently added to `stdpopsim`, which demonstrate some of the main considerations involved in the process of designing realistic chromosome-scale simulations. While the guidelines provided in this paper are intended for any researcher interested in implementing a population genomic simulation using any software, we highlight the ways in which the `stdpopsim` framework eases the burden involved in this process and facilitates reproducible research.

## The utility of stdpopsim for chromosome-scale simulations

We begin by providing a brief overview of the importance of chromosome-scale simulations and the main rationale behind `stdpopsim`; see Adrion et al. (2020) for more on the topic. The main objective of population genomic simulations is to recreate patterns of sequence variation along the genome under the inferred evolutionary history of a given species. To achieve this, `stdpopsim` is built on top of the `msprime` (Kelleher et al., 2016; Nelson et al., 2020; Baumdicker et al., 2021) and `SLiM` (Haller and Messer, 2019) simulation engines, which are capable of producing fairly realistic patterns of sequence variation if provided with accurate descriptions of the genome architecture and evolutionary history of the simulated species. The required parameters include the number of chromosomes and their lengths, mutation and recombination rates, the demographic history of the simulated population, and, potentially, the landscape of natural selection along the genome. A key challenge when setting up a population genomic simulation is to obtain estimates of all of these quantities from the literature and then correctly implement them in an appropriate simulation engine. Detailed estimates of all of these quantities are increasingly available due to the growing availability of population genomic data coupled with methodological advances. Incorporating this data into a population genomic simulation often involves integrating this data between different literature sources, which can require specialized knowledge of population genetics theory. Thus, the process of coding a realistic simulation can be quite time-consuming and often error-prone.

The main objective of `stdpopsim` is to streamline this process, and to make it more robust and more reproducible. Contributors collect parameter values for their species of interest from the literature, and then specify these parameters in a template file for the new model. This model then undergoes a peerreview process, which involves another researcher independently recreating the model based on the provided documentation. Automated scripts then execute to compare the two models; if discrepancies are found in this process, they are resolved by discussion between the contributor and reviewer, and if necessary with input of additional members of the community. This quality-control process quite often finds subtle bugs (e.g., as in Ragsdale et al., 2020) or highlights parts of the model that are ambiguously defined by the literature sources. This increases the reliability and reproducibility of the resulting simulations in any downstream analysis.

Another important goal of `stdpopsim` is to promote and facilitate chromosome-scale simulations, as opposed to the common practice of simulating many short segments (see, e.g., Harris and Nielsen, 2016). Simulation of long sequences, on the order of 10^7^ bases, has until recently been computationally prohibitive, but this has changed with the development of modern simulation engines such as msprime and SLiM. Generating chromosome-scale simulations has several key benefits. First, the organization of genes on chromosomes is a key feature of a species’ genome that is ignored in many traditional population genomic simulations (see Schrider (2020) for one exception). Second, modeling physical linkage allows simulations to capture important correlations between genetic variants on a chromosome. These correlations reduce variance relative to separate and independent simulations of equivalent genetic material. This has a particularly striking effect in long stretches with a low recombination rate, as observed for instance on the long arm of human chromosome 22 (Dawson et al., 2002). In bacteria, a similar effect occurs due to genome-wide linkage that is broken only by horizontal transfer of short segments (Didelot and Maiden, 2010). When conducting simulations with natural selection, linkage has an even stronger effect. Selection acting on a small number of sites can indirectly influence levels and patterns of genetic variation at linked neutral sites, which has been shown to have a widespread effect on patterns of genomic variation in a myriad of species (e.g., McVicker et al., 2009; Charlesworth, 2012). In addition, the lengths of chromosome-scale shared haplotypes within and between populations provides valuable information on their demographic history. Demographic inference methods that use such information, such as MSMC (Schiffels and Wang, 2020) and IBDNe (Browning and Browning, 2015), perform best on long genomic segments with realistic recombination rates. Chromosome-scale simulations are clearly required to test (or train) such methods, or to conduct power analyses when designing empirical studies that use them. With `stdpopsim`, such simulations are available with just a single call to a command-line script or with execution of a handful of lines of Python code.

## Additions to stdpopsim

When first published, the `stdpopsim` catalog included six species: *Homo sapiens, Pongo abelii, Canis familiaris, Drosophila melanogaster, Arabidopsis thaliana*, and *Escherichia coli* (Figure 1). One way the catalog has expanded is through the introduction of additional demographic models for *Homo sapiens*, *Pongo abelii*, *Drosophila melanogaster*, and *Arabidopsis thaliana*, enabling a wider variety of simulations for these well-studied species. However, the initial collection of six species represents only a small slice of the tree of life. This is a concern not only because there is a large community of researchers studying other organisms, but also because methods developed for application to model species (such as humans) may not perform well when applied to other species with very different biology. Adding species to the `stdpopsim` catalog will allow developers to easily test their methods across a wider variety of organisms.

**Figure 1:**
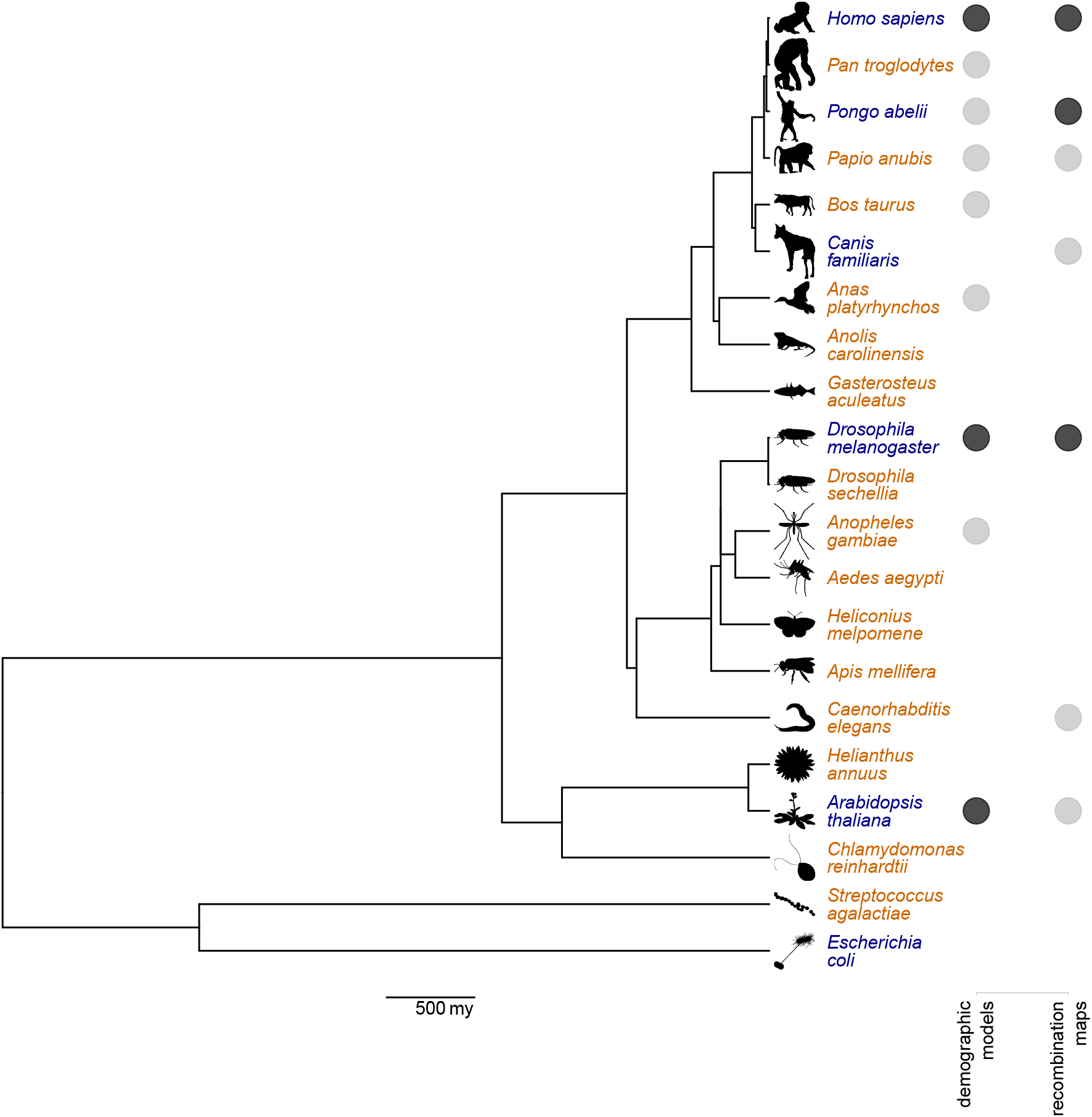
Phylogenetic tree of species available in the `stdpopsim` catalog, including the six species we published in the original release (Adrion et al., 2020, in blue), and 15 species that have since been added (in orange). Solid circles indicate species that have one (light grey) or more (dark grey) demographic models and recombination maps. Branch lengths were derived from the divergence times provided by TimeTree5 (Kumar et al., 2022). The horizontal bar below the tree indicates 500 million years (my).

We thus made a concerted effort to recruit members of the population and evolutionary genetics community to add their species of interest to the `stdpopsim` catalog. This effort involved a series of workshops to introduce potential contributors to `stdpopsim`, followed by a “Growing the Zoo” hackathon organized alongside the 2021 ProbGen conference. The seven initial workshops allowed us to reach a broad community of more than 150 researchers, many of whom expressed interest in adding non-model species to `stdpopsim`. The hackathon was then structured based on feedback from these participants. One month before the hackathon, we organized a final workshop to prepare interested participants, by introducing them to the process of developing a new species model and adding it to the `stdpopsim` code base. Roughly 20 scientists participated in the hackathon (most of whom are included as authors on this paper), which resulted in the addition of 15 species to the `stdpopsim` catalog (Figure 1). The catalog now includes a teleost fish (*Gasterosteus aculeatus*), a bird (*Anas platyrhynchos*), a reptile (*Anolis carolinensis*), a livestock species (*Bos taurus*), six insects including two vectors of human disease (*Aedes aegypti* and *Anopheles gambiae*), a nematode (*Caenorhabditis elegans*), two flowering plants including a crop (*Helianthus annuus*), an algae (*Chlamydomonas reinhardtii*), two bacteria, four primates, and a common mammalian associate of humans (*Canis familiaris*). Not all of these have recombination maps or demographic models (see Figure 1), but this lays a framework for future contributions.

Expanding the species catalog required adding several capabilities to the simulation framework of `stdpopsim`. Some features were added by upgrading the neutral simulation engine, `msprime`, from version 0.7.4 to version 1.0 (Baumdicker et al., 2021). Among other features, this upgrade includes a discrete-site model of mutation, which enables simulating sites with multiple mutations and possibly more than two alleles. Another key feature added to `stdpopsim`’s simulation framework was the ability to model non-crossover recombination. In bacteria and archaea, genetic material can be exchanged through horizontal gene transfer, which can add new genetic material (e.g., via the transfer of plasmids) or replace homologous sequences through homologous recombination (Thomas and Nielsen, 2005; Didelot and Maiden, 2010; Gophna and Altman-Price, 2022). However, the initial version of `stdpopsim` used crossover recombination to stand in for these processes. Although we cannot currently simulate varying gene content (as would be required to simulate the addition of new genetic material by horizontal gene transfer), the `msprime` and `SLiM` simulation engines now allow gene conversion, which has the same effect as non-crossover homologous recombination. Following Cury et al. (2022), we use this to include non-crossover homologous recombination in bacterial and archaeal species. This is done in `stdpopsim` by setting a flag in the species model to indicate that recombination should be modeled without crossovers, and specifying an average tract length of exchanged genetic material. For example, the model for *Escherichia coli* has been updated in the `stdpopsim` catalog to use non-crossover recombination at an average rate of 8.9 × 10^-11^ recombination events per base per generation, with an average tract length of 542 bases (Wielgoss et al., 2011; Didelot et al., 2012). Note that this rate (8.9 × 10^-11^) corresponds to the rate of initiation of a recombined tract.

Recombination without crossover is also prevalent in sexually reproducing species, where it is termed *gene conversion*. Gene conversion affects shorter segments than crossover recombination and creates distinct patterns of genetic diversity along the genome (Korunes and Noor, 2017). Indeed, gene conversion rates in some species are estimated to occur at similar or even higher rates than crossover recombination (Gay et al., 2007; Comeron et al., 2012; Wijnker et al., 2013). To accommodate this in `stdpopsim` simulations, one needs to specify the fraction of recombinations that occur due to gene conversion (i.e., without crossover), and the average tract length. For example, the model for *Drosophila melanogaster* has been updated in the `stdpopsim` catalog to have a fraction of gene conversions of 0.83 (in all chromosomes that undergo recombination) and an average tract length of 518 bases (Comeron et al., 2012). This update does not affect the rate of crossover recombination, but it adds gene conversion events at a ratio of 83:17 relative to crossover recombination events. We note that since non-crossover recombination incurs a high computation load in simulation, it is turned off by default in `stdpopsim`, and must be explicitly invoked by the simulation model. Note that ignoring gene conversion may result in a slightly skewed distribution of shared haplotypes between individuals (see Table 1).

**Table 1:** Guidelines for dealing with missing parameters. For each parameter, we provide a suggested course of action, and mention the main discrepancies between simulated data and real genomic data that could be caused by misspecification of that parameter.

| Missing parameter | Suggested action | Possible discrepancies |
| --- | --- | --- |
| Mutation rate | Borrow from closest relative with a citable mutation rate | Number of polymorphic sites |
| Recombination rate | Borrow from closest relative with a citable recombination rate | Patterns of linkage disequilibrium |
| Gene conversion rate and tract length | Set rate to 0 or borrow from closest relative with a citable rate | Lengths of shared haplotypes across individuals |
| Demographic model | Set the effective population size ($N_e$) to a value that reflects the observed genetic diversity | Features of genetic diversity that are captured by the site frequency spectrum, such as the prevalence of low-frequency alleles |

Another important extension of `stdpopsim` allows augmenting a genome assembly with genome annotations, such as coding regions, promoters, and conserved elements. These annotations can be used to simulate selection at a subset of sites (such as the annotated coding regions) using parametric distributions of fitness effects. Standardized, easily accessible simulations that include the reality of pervasive linked selection in a species-specific manner has long been identified as a goal for evolutionary genetics (e.g., McVicker et al., 2009; Comeron, 2014). Thus, we expect this extension of `stdpopsim` to be transformative in the way simulations are carried out in population genetics. This significant new capability of the `stdpopsim` library will be detailed in a forthcoming publication, and is not the focus of this paper.

## Guidelines for implementing a population genomic simulation

The concentrated effort to add species to the `stdpopsim` catalog has led to a series of important insights about this process, which we summarize here as a set of guidelines for implementing realistic simulations for any species. Our intention is to provide general guidance that applies to any population genomic simulation software, but we also mention specific requirements that apply to simulations done in `stdpopsim`.

### Basic setup for chromosome-level simulations

Implementing a realistic population genomic simulation for a species of interest requires a detailed description of the organism’s demography and mechanisms of genetic inheritance. While simulation software requires unforgivingly precise values, in practice we may only have rough guesses for most of the parameters describing these processes. In this section, we list the relevant parameters and provide guidelines for how to set them based on current knowledge.

1. **A chromosome-level genome assembly**, which consists of a list of chromosomes or scaffolds and their lengths. Having a good quality assembly with complete chromosomes, or at least very long scaffolds, is necessary if chromosome-level population genomic simulations are to reflect the genomic architecture of the species. When expanding the `stdpopsim` catalog during the “Growing the Zoo” hackathon, we considered the possibility of adding species whose genome assemblies are composed of many relatively small contigs, unanchored to chromosome-level scaffolds. Although we had not previously put restrictions on which species might be added, we decided that we would only add species with chromosome-level assemblies. The main justification for this restriction is that species with less complete genome builds typically do not have good recombination maps and demographic models, making chromosome-level simulation much less useful in such species. Another issue is the storage burden and long load times involved in dealing with hundreds of contigs. Finally, each species requires validation of its code before it is added to the `stdpopsim` catalog, as well as long-term maintenance to keep it up-to-date with changes made to the `stdpopsim` framework. So, the benefit of including species with very partial genome builds in `stdpopsim` would be outweighed by the substantial extra burden on `stdpopsim` maintainers as well as downstream users of these models. Another reason to focus on species with chromosome-level assemblies is that we expect their numbers to dramatically increase in the near future due to numerous genome initiatives (Lewin et al., 2022; Rhie et al., 2021; Cheng et al., 2018) and the development of new long-read sequencing technologies and assembly pipelines (Chakraborty et al., 2016; Amarasinghe et al., 2020, 2021).
2. **An average mutation rate** for each chromosome (per generation per bp). This rate estimate can be based on sequence data from pedigrees, mutation accumulation studies, or comparative genomic analysis calibrated by fossil data (i.e., phylogenetic estimates). At present, `stdpopsim` simulates mutations at a constant rate under the Jukes–Cantor model of nucleotide mutations (Jukes and Cantor, 1969). However, we anticipate future development will provide support for more complex, heterogeneous mutational processes, as these are easily specified in both the `SLiM` and `msprime` simulation engines. Such progress will further improve the realism of simulated genomes, since mutation processes, including rates, are known to vary along the genome and through time (Benzer, 1961; Ellegren et al., 2003; Supek and Lehner, 2019).
3. **Recombination rates** (per generation per bp). Ideally, a population genomic simulation should make use of a chromosome-level recombination map, since the recombination rate is known to vary widely across chromosomes (Nachman, 2002), and this can strongly affect the patterns of linkage disequilibrium and shared haplotype lengths. When this information is not available, we suggest specifying an average recombination rate for each chromosome. At minimum, an average genomewide recombination rate needs to be specified, which is typically available for well-assembled genomes. For bacteria and archaea, which primarily experience non-crossover recombination, the average tract length should also be specified (see details in previous section). **Gene conversion (optional):** If one wishes to model gene conversion in eukaryotes, either together with crossover recombination or as a stand-alone process, then one should specify the fraction of recombinations done by gene conversion as well as the per chromosome average tract length.
4. **A demographic model** describing ancestral population sizes, split times, and migration rates. Selection of a reasonable demographic model is often crucial, since misspecification of the model can generate unrealistic patterns of genetic variation that will affect downstream analyses (e.g., Navascués and Emerson, 2009). A given species might have more than one demographic model, fit from different data or by different methods. Thus, when selecting a demographic model, one should examine the data sources and methods used to obtain it to ensure that they are relevant to the study at hand (see also **Limiations of simulated genomes** below). At a minimum, simulation requires a single estimate of effective population size. This estimate, which may correspond to some sort of historical average effective population size, should produce simulated data that matches the average observed genetic diversity in that species. Note, however, that this average effective population size cannot capture features of genetic variation that are caused by recent changes in population size and the presence of population structure (MacLeod et al., 2013; Eldon et al., 2015). For example, a recent population expansion will produce an excess of low-frequency alleles that no simulation of a constant-sized population will reproduce (Tennessen et al., 2012).
5. **An average generation time** for the species. This parameter is an important part of the species’ natural history. This value does not directly affect the simulation, since `stdpopsim` uses either the Wright–Fisher model (in `SLiM`) or the Moran model (in `msprime`), both of which operate in time units of generations. Thus, the average generation time is only currently used to convert time units to years, which is useful when comparing among different demographic models.

These five categories of parameters are sufficient for generating simulations under neutral evolution. Such simulations are useful for a number of purposes, but they cannot be used to model the influence of natural selection on patterns of genetic variation. To achieve this, the simulator needs to know which regions along the genome are subject to selection, and the nature and strength of this selection. As mentioned above, the ability to simulate chromosomes with realistic models of selection is still under development, and will be finalized in the next release of `stdpopsim`. The development version of `stdpopsim` enables simulation with selection (using the `SLiM` engine) by specifying genome annotations and distributions of fitness effects, as specified below.

6. **Genome annotations**, specifying regions subject to selection (as, for example, a GFF3/GTF file). For instance, annotations can contain information on the location of coding regions, the position of specific genes, or conserved non-coding regions. Regions not covered by the annotation file are assumed to be evolving free from the effects of direct natural selection.
7. **Distributions of fitness effects** (DFEs) for each annotation. Each annotation is associated with a DFE describing the probability distribution of selection coefficients (deleterious, neutral, and beneficial) for mutations occurring in the region covered by the annotation. DFEs can be inferred from population genomic data (reviewed in Eyre-Walker and Keightley, 2007), and are available for several species (e.g., Ma et al., 2013; Huber et al., 2018).

The current release of `stdpopsim` contains annotations and implemented DFE models for the three model species: *A. thaliana*, *D. melanogaster*, and *H. sapiens*. A forthcoming publication will provide details about how this is implemented in `stdpopsim` and examples of possible uses of this feature.

### Extracting parameters from the literature

Simulations cannot of course precisely match reality, but in setting up simulations it is desirable to choose parameters that best reflect our current understanding of the evolutionary history of the species of interest. In practice a researcher may choose each parameter to match a fairly precise estimate or a wild guess, which may be obtained from a peer-reviewed publication or by word of mouth. However, values in `stdpopsim` are always chosen to match published estimates, so that the underlying data and methods are documented and can be validated. Because the process of converting information reported in the literature to parameters used by a simulation engine is quite error-prone, independent validation of the simulation code is crucial. We highly recommend following a quality-control procedure similar to the one used in `stdpopsim`, in which each species or model added to the catalog is independently recreated or thoroughly reviewed by a separate researcher.

Obtaining reliable and citable estimates for all model parameters is not a trivial task. Oftentimes, values for different parameters must be gleaned from multiple publications and combined. For example, it is not uncommon to find an estimate of a mutation rate in one paper, a recombination map in a separate paper, and a suitable demographic model in a third paper. Integrating information from different publications requires caution, since some of these parameter estimates are entangled in non-trivial ways. For instance, consider simulating a demographic model estimated in a specific paper that assumes a certain mutation rate. Naively using the demographic model, as published, with a new estimate of the mutation rate will lead to levels of genetic diversity that do not fit the genomic data. This is addressed in `stdpopsim` by allowing a demographic model to be simulated using a mutation rate that differs from the default rate specified for the species. See, for example, the model implemented for *Bos taurus*, which is described in the next section. This important feature does not necessarily fix all potential inconsistencies caused by assumptions made by the demographic inference method (such as assumptions about recombination rates). It is therefore recommended, when possible, to take the demographic model, mutation rates, and recombination rates from the same study, and to proceed carefully when mixing sources. An additional tricky source of inconsistency is coordinate drift between subsequent versions of genome assemblies. In `stdpopsim`, we follow the UCSC Genome Browser and use liftOver to convert the coordinates of recombination maps and genome annotations to the coordinates of the current genome assembly (Hinrichs et al., 2006).

### Limitations of simulated genomes

Despite their great utility, simulated genomes cannot fully capture all aspects of genetic variation as observed in real data, with some aspects modeled better than others. As mentioned above, this will strongly depend on the demographic model used in simulation. Thus, it is important to consider potential limitations of different demographic models in reflecting observed genetic variation. First, a demographic model inferred from analysis of genomic data will likely depend on the samples that contributed the analyzed genomes. The inferred demographic model can only reflect the genealogical ancestry of these sampled individuals, and this will typically make up a small portion of the complete genealogical ancestry of the species. Thus, demographic models inferred from larger sets of samples from diverse ancestry backgrounds may potentially provide a more comprehensive depiction of genetic variation within a species. This is true if sufficiently realistic demographic models can be fit—models that account for the structure of populations within a species. That said, the choice of samples used for inference will mostly influence recent changes in genetic variation. This is because the genealogy of even a single individual consists of numerous ancestors in each generation in the deep past, which is the premise of methods that infer ancestral population sizes from a single input genome (Li and Durbin, 2011).

The computational method used for inference also affects the way genetic variation is reflected by the demographic model, because different methods derive their inference from different features of genomic variation. Some methods make use of the site frequency spectrum at unlinked single sites (e.g., Gutenkunst et al., 2009; Excoffier et al., 2013; Liu and Fu, 2015), while other methods use haplotype structure (e.g., Li and Durbin, 2011; Schiffels and Wang, 2020; Browning and Browning, 2015). This, in turn, may influence the accuracy of different features in the inferred demography. For example, very recent demographic changes, such as recent admixture or bottlenecks, are difficult to infer from the site frequency spectrum, but are more easily inferred by examining shared long haplotypes (as demonstrated by the demographic model inferred for *Bos taurus* by MacLeod et al. (2013); see below). Several studies have compared different approaches to demographic inference (e.g., Harris and Nielsen, 2013; Beichman et al., 2017), but unfortunately, there is currently no succinct handbook that describes the relative strengths and weaknesses of different methods. Thus, assessing the potential limitations of a given demographic model currently requires some familiarity with the method used for its inference. In addition, all methods assume that the input sequences are neutrally evolving. This implies that technical choices, such as the specific genomic segments analyzed and various filters, may also influence the inferred model and its ability to model observed genetic variation. Thus, it is strongly advised to read the study that inferred the demographic model and understand potential limitations that stem from the selection of samples, methods, and filters.

We note that inclusion of a demographic model in the `stdpopsim` catalog does not involve any judgment as to which aspects of genetic variation it captures. Any model that is a faithful implementation of a published model inferred from genomic data can be added to the `stdpopsim` catalog. Thus, potential users of `stdpopsim` should use the implemented models with the appropriate caution, keeping in mind the limitations discussed above. We maintain a fairly detailed documentation page for the catalog (see **Data availability**), which contains a brief summary for each demographic model. This summary includes a graphical description of the model (such as the one shown for *Anopheles gambiae* in Fig. 2B), as well as a description of the data and method used for inference. Therefore, the documentation can provide guidance to potential users of `stdpopsim` in the process of selecting an appropriate demographic model for simulation. Finally, we hope that the standardized simulations implemented in `stdpopsim` will facilitate additional studies that examine the relative strengths and limitations of different approaches to demographic inference (and modeling genetic variation in general), and this will allow us to generate even more realistic simulations in the future.

**Figure 2:**
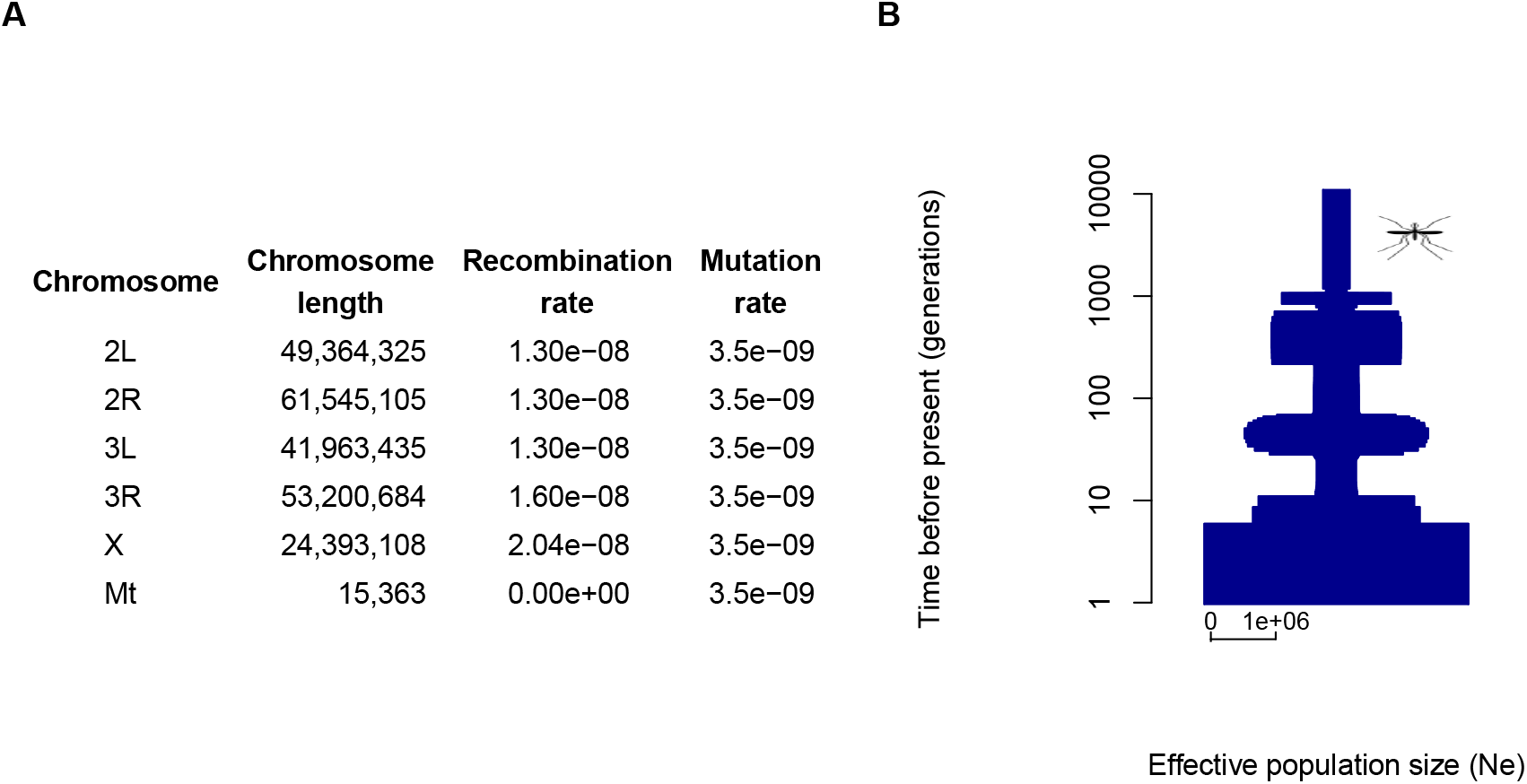
The species parameters and demographic model used for *Anopheles gambiae* in the `stdpopsim` catalog. (A) The parameters associated with the genome build and species, including chromosome lengths, average recombination rates (per base per generation), and average mutation rates (per base per generation). (B) A graphical depiction of the demographic model, which consists of a single population whose size changes throughout the past 11,260 generations in 67 time intervals (note the log scale). The width at each point depicts the effective population size (*N_e_*), with the horizontal bar at the bottom indicating the scale for *N_e_* = 10^6^. This figure is adapted from the data on the `stdpopsim` catalog documentation page (see **Data availability**) and plotted with POPdemog (Zhou et al., 2018).

### Filling in the missing pieces

For many species it is difficult to obtain estimates of all necessary model parameters. Table 1 provides suggestions for ways to deal with missing values of various model parameters. The table also mentions possible consequences of misspecification of each parameter.

In some cases, one may wish to generate simulations for a species with a partial genome build. Despite the focus of `stdpopsim` on species with chromosome-level assemblies (see discussion above), simulation is still potentially useful for species with less complete assemblies, with some important considerations to keep in mind. Longer contigs or scaffolds in these builds can be simulated separately and independently. This approach allows us to model genetic linkage within each contig, but linkage between different contigs that map to the same chromosome will not be captured by the simulation. This provides a reasonable approximation for many purposes, at least for genomic regions far from the contig edges. For shorter contigs, separate independent simulations will not be able to capture patterns of long-range linkage in a reasonably realistic way. Thus, a potentially viable option for shorter contigs is to combine them into longer pseudochromosomes, trying to mimic the species’ expected chromosome lengths. Despite their somewhat artificial construction, these pseudo-chromosomes have the important benefit of capturing patterns of linkage similar to those observed in real genomic chromosomes. If, for example, the main purpose of the simulation is to examine the distribution of lengths of shared haplotypes between individuals, or study patterns of background selection, then it makes sense to simulate such pseudo-chromosomes. However, genetic correlations between different specific contigs lumped together in this way are obviously not accurate. So, if the main purpose of the simulation is to examine local patterns of genetic variation in loci of interest, then it may be more appropriate to simulate the relevant contigs separately (even if they are short), or to randomly sample several mappings of contigs to pseudo-chromosomes. For some purposes it makes sense to simulate a large number of unlinked sites (Gutenkunst et al., 2009; Excoffier et al., 2013), which can be generated without any sort of genome assembly. However, this approach would not have the benefits of chromosome-scale simulations. While some of the same considerations hold when simulating unlinked short sequences, a detailed discussion about such simulations goes beyond the scope of this paper. Ultimately, the recommended mode of simulation for a species with a partial genome assembly depends on the intended use of the simulated genomes.

### Examples of added species

In this section, we provide examples of two species recently added to the `stdpopsim` catalog, *Anopheles gambiae* and *Bos taurus*, to demonstrate some of the key considerations of the process. In each example, we highlight in bold the model parameters set for each species.

#### *Anopheles gambiae* (mosquito)

*Anopheles gambiae*, the African malaria mosquito, is a non-model organism whose population history has direct implications for human health. Several large-scale studies in recent years have provided information about the population history of this species on which population genomic simulations can be based (e.g., Miles et al., 2017; Clarkson et al., 2020). The genome assembly structure used in the species model is from the AgamP4 **genome assembly** (Sharakhova et al., 2007), downloaded directly from Ensembl (Howe et al., 2020) using the special utilities provided by `stdpopsim`.

Estimates of average **recombination rates** for each of the chromosomes (excluding the mitochondrial genome) were taken from a recombination map inferred by Pombi et al. (2006) which itself included information from Zheng et al. (1996) (Figure 2A). As direct estimates of **mutation rate** (e.g., via mutation accumulation) do not currently exist for *Anopheles gambiae*, we used the genome-wide average mutation rate of *μ* = 3.5 × 10^-9^ mutations per generation per site estimated by Keightley et al. (2009) for the fellow Dipteran *Drosophila melanogaster*, a rate that was used for analysis of *A. gambiae* data in Miles et al. (2017). To obtain an estimate for the default **effective population size** (N_e_), we used the formula *θ* = 4*μN_e_*, with the above mutation rate (μ = 3.5 × 10^-9^ mutations per base per generation) and a mean nucleotide diversity of θ ≈ 0.015, as reported by Miles et al. (2017) for the Gabon population. This resulted in an estimate of N_e_ = 1.07 × 10^6^, which we rounded down to one million. These steps were documented in the code for the `stdpopsim` species model, to facilitate validation and future updates. We acknowledge that some of these steps involve somewhat arbitrary choices, such as the choice of the Gabon population and rounding down of the final value. However, this should not be seen as a considerable source of misspecification, since this value of N_e_ is meant to provide only a rough approximation to historical population sizes and would be overwritten by a more detailed demographic model. Miles et al. (2017) inferred demographic models from *Anopheles* samples from nine different populations (locations) using the stairway plot method (Liu and Fu, 2015). We chose to include in `stdpopsim` the **demographic model** inferred from the Gabon sample, which consists of a single population whose size fluctuated from below 80,000 (an ancient bottleneck roughly 10,000 generations ago) to the present-day estimate of over 4 million individuals (Figure 2B). To convert the timescale from generations to years, we used an **average generation time** of 1/11 years, as in Miles et al. (2017).

All of these parameters were set in the species entry in the `stdpopsim` catalog, accompanied by the relevant citation information, and the model underwent the standard quality-control process. The species entry may be refined in the future by adding more demographic models, updating or refining the recombination map, or updating the mutation rate estimates based on ones directly estimated for this species. Note that even if the mutation rate is updated sometime in the future, the demographic model mentioned above should still be associated with the current mutation rate (μ = 3.5 × 10^-9^ mutations per base per generation), since this was the rate used in its inference.

#### *Bos taurus* (cattle)

*Bos taurus* (cattle) was added to the `stdpopsim` catalog during the 2020 hackathon because of its agricultural importance. Agricultural species experience strong selection due to domestication and selective breeding, leading to a reduction in effective population size. These processes, as well as admixture and introgression, produce patterns of genetic variation that can be very different from typical model species (Larson and Burger, 2013). These processes have occurred over a relatively short period of time, since the advent of agriculture roughly 10,000 years ago, and they have intensified over the years to improve food production (Gaut et al., 2018; MacLeod et al., 2013). High-quality genome assemblies are now available for several breeds of cattle (e.g., Rosen et al., 2020; Heaton et al., 2021; Talenti et al., 2022) and the use of genomic data has become ubiquitous in selective breeding (Meuwissen et al., 2001; MacLeod et al., 2014; Obšteter et al., 2021; Cesarani et al., 2022). Modern cattle have extremely low and declining genetic diversity, with estimates of effective population size around 90 in the early 1980s (MacLeod et al., 2013; VanRaden, 2020; Makanjuola et al., 2020). On the other hand, the ancestral effective population size is estimated to be roughly N_e_=62,000 (MacLeod et al., 2013). This change in effective population size presents a challenge for demographic inference, selection scans, genome-wide association, and genomic prediction (MacLeod et al., 2013, 2014; Hartfield et al., 2022). For these reasons, it was useful to develop a detailed simulation model for cattle to be added to the `stdpopsim` catalog.

We used the most recent **genome assembly**, ARS-UCD1.2 (Rosen et al., 2020), a constant **mutation rate** of μ = 1.2 × 10^-8^ mutations per base per generation for all chromosomes (Harland et al., 2017), and a constant **recombination rate** of *r* = 9.26 × 10^-9^ recombinations per base per generation for all chromosomes other than the mitochondrial genome (Ma et al., 2015). With respect to the **effective population size**, it is clear that simulating with either the ancestral or current effective population size would not generate realistic genome structure and diversity (MacLeod et al., 2013; Rosen et al., 2020). Since `stdpopsim` does not allow for a missing value of *N_e_*, we chose to set the species default *N_e_* to the ancestral estimate of 6.2 × 10^4^. However, we strongly caution that simulating the cattle genome with any fixed value for N_e_ will generate unrealistic patterns of genetic variation, and recommend using a reasonably detailed demographic model. Note that the default N_e_ is only used in simulation if a demographic model is not specified. To this end, we implemented the **demographic model** of the Holstein breed, which was inferred by MacLeod et al. (2013) from runs of homozygosity in the whole-genome sequence of two iconic bulls. This demographic model specifies changes in the ancestral effective population size from N_e_=62,000 at around 33,000 generations ago to N_e_=90 in the 1980s in a series of 13 instantaneous population size changes (taken from Supplementary Table S1 in MacLeod et al., 2013). To convert the timescale from generations to years, we used an **average generation time** of 5 years (MacLeod et al., 2013). Note that this demographic model does not capture the intense selective breeding since the 1980s that has even further reduced the effective population size of cattle (MacLeod et al., 2013; VanRaden, 2020; Makanjuola et al., 2020). These effects can be modeled with downstream breeding simulations (e.g., Gaynor et al., 2020).

When setting up the parameters of the demographic model, we noticed that the inference by MacLeod et al. (2013) assumed a genome-wide fixed recombination rate of *r* = 10^-8^ recombinations per base per generation, and a fixed mutation rate of *μ* = 9.4 × 10^-9^ mutations per base per generation (considering also sequence errors). The more recently updated mutation rate assumed in the species model (1.2 × 10^-8^ mutations per base per generation, from Harland et al., 2017) is thus 28% higher than the rate used for inference. As a result, if genomes were simulated under this demographic model with the species’ default mutation rate they would have considerably higher sequence diversity than actually observed in real genomic data. To address this, we specified a mutation rate of *μ* = 9.4 × 10^-9^ in the demographic model, which then overrides the species’ mutation rate when this demographic model is applied in simulation. The issue of fitting the rates used in simulation with those assumed during inference was discussed during the independent review of this demographic model, and it raised an important question about recombination rates. Since MacLeod et al. (2013) use runs of homozygosity to infer the demographic model, their results depends on the assumed recombination rate. The recombination rate assumed in inference (*r* = 10^-8^ recombinations per base per generation) is 8% higher than the one used in the species model (*r* = 9.26 × 10^-9^). In its current version, `stdpopsim` does not allow specification of a separate recombination rate for each demographic model, so we had no simple way to adjust for this. Future versions of `stdpopsim` will enable such flexibility. Thus, we note that genomes simulated under this demographic model as currently implemented in `stdpopsim` might have slightly higher linkage disequilibrium than observed in real cattle genomes. However, we anticipate that this would affect patterns less than selection due to domestication and selective breeding, which are not yet modeled at all in `stdpopsim` simulations.

## Conclusion

As our ability to sequence genomes continues to advance, the need for population genomic simulations of new model and non-model organisms is becoming acute. So, too, is the concomitant need for an expandable framework for implementing such simulations and guidance for how to do so. Generating realistic wholegenome simulations presents significant challenges both in coding and in choosing parameter values on which to base the simulation. With `stdpopsim`, we provide a resource that is uniquely poised to address these challenges as it provides easy access to state-of-the-art simulation engines and practices, and an easy procedure for including new species. Moreover, we aim for the choices regarding inclusion of new species to be driven by the needs of the population genomics community. In this manuscript we describe the expansion of `stdpopsim` in two ways: the addition of new features to the simulation framework that incorporate new evolutionary processes, such as non-crossover recombination, broadening the diversity of species that can be realistically modeled; and the considerable expansion of the catalog itself to include more species and demographic models.

We also formulated a series of guidelines for implementing population genomic simulations, based on insights from the community-driven process of expanding the `stdpopsim` catalog. These guidelines specify the basic requirements for generating a useful chromosome-level simulation for a given species, as well as the rationale behind these requirements. We also discuss special considerations for collecting relevant information from the literature, and what to do if some of that information is not available. Because this process is quite error-prone, we encourage wider adoption of “code review”: researchers implementing simulations should have their parameter choices and implementation reviewed by at least one other researcher. The guidelines in this paper can be followed when implementing a simulation independently for a single study, or (as we encourage others to do) when adding code to `stdpopsim`, which helps to ensure its robustness and to make it available for future research. Currently, large-scale efforts such as the Earth Biogenome and its affiliated project networks are generating tens of thousands of genome assemblies. Each of these assemblies would become a candidate for inclusion into the `stdpopsim` catalog, although substantial changes to the structure of `stdpopsim` would be required to include so many distinct species. As annotations of those genome assemblies improve over time, this information, too, can easily be added to the `stdpopsim` catalog.

One of the important objectives of the PopSim consortium is to leverage `stdpopsim` as a means to promote education and inclusion of new communities into computational biology and software development. We are keen to use outreach, such as the workshops and hackathons described here, as a way to grow the `stdpopsim` catalog and library while also democratizing the development of population genomic simulations in general. We predict that the increased use of chromosome-scale simulations in non-model species will lead to an improvement in inference methods, which traditionally have been quite narrowly focused on well-studied model organisms. Thus, we hope that further expansion of `stdpopsim` will improve the ease and reproducibility of research across a larger number of systems, while simultaneously expanding the community of software developers among population and evolutionary geneticists.

## Data availability

The code for `stdpopsim` and the species catalog are available from: https://github.com/popsim-consortium/stdpopsim. The documentation page for the `stdpopsim` catalog is available from: https://popsim-consortium.github.io/stdpopsim-docs/stable/catalog.html

## Acknowledgments

We wish to thank the dozens of workshop attendees, and especially the two dozen or so hackathon participants, whose combined feedback motivated many of the updates made to `stdpopsim` in the past two years.

## References

Jeffrey R Adrion, Christopher B Cole, Noah Dukler, Jared G Galloway, Ariella L Gladstein, Graham Gower, Christopher C Kyriazis, Aaron P Ragsdale, Georgia Tsambos, Franz Baumdicker, Jedidiah Carlson, Reed A Cartwright, Arun Durvasula, Ilan Gronau, Bernard Y Kim, Patrick McKenzie, Philipp W Messer, Ekaterina Noskova, Diego Ortega-Del Vecchyo, Fernando Racimo, Travis J Struck, Simon Gravel, Ryan N Gutenkunst, Kirk E Lohmueller, Peter L Ralph, Daniel R Schrider, Adam Siepel, Jerome Kelleher, and Andrew D Kern. A community-maintained standard library of population genetic models. eLife, 9:e54967, jun 2020. ISSN 2050-084X. doi: 10.7554/eLife.54967. URL https://doi.org/10.7554/eLife.54967.

Shanika L. Amarasinghe, Shian Su, Xueyi Dong, Luke Zappia, Matthew E. Ritchie, and Quentin Gouil. Opportunities and challenges in long-read sequencing data analysis. Genome Biology, 21, 2020. doi: https://doi.org/10.1186/s13059-020-1935-5.

Shanika L Amarasinghe, Matthew E Ritchie, and Quentin Gouil. long-read-tools.org: an interactive catalogue of analysis methods for long-read sequencing data. GigaScience, 10(2), 02 2021. ISSN 2047-217X. doi: 10.1093/gigascience/giab003. URL https://doi.org/10.1093/gigascience/giab003.giab003.

Franz Baumdicker, Gertjan Bisschop, Daniel Goldstein, Graham Gower, Aaron P Ragsdale, Georgia Tsambos, Sha Zhu, Bjarki Eldon, E Castedo Ellerman, Jared G Galloway, Ariella L Gladstein, Gregor Gorjanc, Bing Guo, Ben Jeffery, Warren W Kretzschumar, Konrad Lohse, Michael Matschiner, Dominic Nelson, Nathaniel S Pope, Consuelo D Quinto-Cortés, Murillo F Rodrigues, Kumar Saunack, Thibaut Sellinger, Kevin Thornton, Hugo van Kemenade, Anthony W Wohns, Yan Wong, Simon Gravel, Andrew D Kern, Jere Koskela, Peter L Ralph, and Jerome Kelleher. Efficient ancestry and mutation simulation with msprime 1.0. Genetics, 220(3), 12 2021. ISSN 1943-2631. doi: 10.1093/genetics/iyab229. URL https://doi.org/10.1093/genetics/iyab229.iyab229.

A. C. Beichman, T. N. Phung, and K. E. Lohmueller. Comparison of Single Genome and Allele Frequency Data Reveals Discordant Demographic Histories. G3 (Bethesda), 7(11):3605–3620, Nov 2017.

Annabel C. Beichman, Emilia Huerta-Sanchez, and Kirk E. Lohmueller. Using genomic data to infer historic population dynamics of nonmodel organisms. Annu. Rev. Ecol. Evol. Syst., 49:433–456, 2018. ISSN 15452069. doi: 10.1146/annurev-ecolsys-110617-062431.

Seymour Benzer. On the topography of the genetic fine structure. Proceedings of the National Academy of Sciences, 47(3):403–415, 1961. doi: 10.1073/pnas.47.3.403. URL https://www.pnas.org/doi/abs/10.1073/pnas.47.3.403.

Paul D. Blischak, Michael S. Barker, Ryan N. Gutenkunst, and Daniel Falush. Inferring the demographic history of inbred species from genome-wide SNP frequency data. Mol. Biol. Evol., 37(7):2124–2136, 2020. ISSN 15371719. doi: 10.1093/molbev/msaa042.

Sharon R. Browning and Brian L. Browning. Accurate non-parametric estimation of recent effective population size from segments of identity by descent. The American Journal of Human Genetics, 97 (3):404–418, 2015. ISSN 0002-9297. doi: https://doi.org/10.1016/j.ajhg.2015.07.012. URL https://www.sciencedirect.com/science/article/pii/S0002929715002888.

A Cesarani, D Lourenco, S Tsuruta, A Legarra, E L Nicolazzi, P M VanRaden, and I Misztal. Multibreed genomic evaluation for production traits of dairy cattle in the United States using single-step genomic best linear unbiased predictor. Journal of Dairy Science, 105(6):5141–5152, 2022. doi: https://doi.org/10.3168/jds.2021-21505.

Mahul Chakraborty, James G Baldwin-Brown, Anthony D Long, and JJ Emerson. Contiguous and accurate *de novo* assembly of metazoan genomes with modest long read coverage. Nucleic acids research, 44(19): e147–e147, 2016.

Brian Charlesworth. The effects of deleterious mutations on evolution at linked sites. Genetics, 190(1):5–22, Jan 2012.

Shifeng Cheng, Michael Melkonian, Stephen A. Smith, Samuel Brockington, John M. Archibald, Pierre-Marc Delaux, Fay-Wei Li, Barbara Melkonian, Evgeny V. Mavrodiev, Wenjing Sun, Yuan Fu, Huanming Yang, Douglas E. Soltis, Sean W. Graham, Pamela S. Soltis, Xin Liu, Xun Xu, and Gane Ka-Shu Wong. 10KP: A phylodiverse genome sequencing plan. Gigascience, 3(7), 2018. ISSN 2047-217X. doi: 10.1093/gigascience/giy013.

Chris S Clarkson, Alistair Miles, Nicholas J Harding, Eric R Lucas, CJ Battey, Jorge Edouardo Amaya-Romero, Andrew D Kern, Michael C Fontaine, Martin J Donnelly, Mara KN Lawniczak, et al. Genome variation and population structure among 1142 mosquitoes of the African malaria vector species *Anopheles gambiae* and Anopheles coluzzii. Genome research, 30(10):1533–1546, 2020.

Josep M Comeron. Background selection as baseline for nucleotide variation across the Drosophila genome. PLoS Genetics, 10(6):e1004434, 2014.

Josep M. Comeron, Ramesh Ratnappan, and Samuel Bailin. The many landscapes of recombination in *Drosophila melanogaster*. PLoS Genet, 8(10):e1002905, 2012.

Katalin Csilléry, Michael G B Blum, Oscar E Gaggiotti, and Olivier François. Approximate Bayesian Computation (ABC) in practice. Trends Ecol. Evol., 25(7):410–8, jul 2010. ISSN 0169-5347. doi: 10.1016/j.tree.2010.04.001. URL http://www.ncbi.nlm.nih.gov/pubmed/20488578.

J. Cury, B. C. Haller, G. Achaz, and F. Jay. Simulation of bacterial populations with SLiM. Peer Community Journal, 2:e7, 2022. doi: 10.24072/pcjournal.72. URL https://peercommunityjournal.org/articles/10.24072/pcjournal.72/.

A. D. Cutter and B. A. Payseur. Genomic signatures of selection at linked sites: unifying the disparity among species. Nature Reviews Genetcs, 14(4):262–274, 2013. doi: https://doi.org/10.1038/nrg3425. URL https://www.nature.com/articles/nrg3425.

Darwin Tree of Life Project Consortium. Sequence locally, think globally: The Darwin Tree of Life Project. Proceedings of the National Academy of Sciences, 119(4):e2115642118, 2022.

Elisabeth Dawson, Gonçalo R Abecasis, Suzannah Bumpstead, Yuan Chen, Sarah Hunt, David M Beare, Jagjit Pabial, Thomas Dibling, Emma Tinsley, Susan Kirby, et al. A first-generation linkage disequilibrium map of human chromosome 22. Nature, 418(6897):544–548, 2002.

X. Didelot and M. C. Maiden. Impact of recombination on bacterial evolution. Trends Microbiol, 18(7): 315–322, Jul 2010.

X. Didelot, G. Meric, D. Falush, and A. E. Darling. Impact of homologous and non-homologous recombination in the genomic evolution of *Escherichia coli*. BMC Genomics, 13:256, Jun 2012.

B. Eldon, M. Birkner, J. Blath, and F. Freund. Can the site-frequency spectrum distinguish exponential population growth from multiple-merger coalescents? Genetics, 199(3):841–856, Mar 2015.

Hans Ellegren. Genome sequencing and population genomics in non-model organisms. Trends Ecol. Evol., 29(1):51–63, 2014. ISSN 01695347. doi: 10.1016/j.tree.2013.09.008. URL http://dx.doi.org/10.1016/j.tree.2013.09.008.

Hans Ellegren, Nick GC Smith, and Matthew T Webster. Mutation rate variation in the mammalian genome. Current Opinion in Genetics & Development, 13(6):562–568, 2003. ISSN 0959-437X. doi: https://doi.org/10.1016/j.gde.2003.10.008. URL https://www.sciencedirect.com/science/article/pii/S0959437X03001461.

Laurent Excoffier, Isabelle Dupanloup, Emilia Huerta-Sánchez, Vitor C. Sousa, and Matthieu Foll. Robust demographic inference from genomic and SNP data. PLOS Genetics, 9(10):1–17, 10 2013. doi: 10.1371/journal.pgen.1003905. URL https://doi.org/10.1371/journal.pgen.1003905.

Adam Eyre-Walker and Peter D Keightley. The distribution of fitness effects of new mutations. Nat. Rev. Genet., 8(8):61061–8, 2007. ISSN 1471-0056. doi: 10.1038/nrg2146.

B S Gaut, D K Seymour, Q Liu, and Y Zhou. Demography and its effects on genomic variation in crop domestication. Nature Plants, 2018. doi: 10.1038/s41477-018-0210-1. URL https://doi.org/10.1038/s41477-018-0210-1.

J. Gay, S. Myers, and G. McVean. Estimating meiotic gene conversion rates from population genetic data. Genetics, 177(2):881–894, Oct 2007.

R Chris Gaynor, Gregor Gorjanc, and John M Hickey. AlphaSimR: an R package for breeding program simulations. G3 Genes—Genomes—Genetics, 11(2), 12 2020. ISSN 2160-1836. doi: 10.1093/g3journal/jkaa017. URL https://doi.org/10.1093/g3journal/jkaa017.jkaa017.

U. Gophna and N. Altman-Price. Horizontal Gene Transfer in Archaea-From Mechanisms to Genome Evolution. Annu Rev Microbiol, 76:481–502, Sep 2022.

G. Gower, P. I. Picazo, M. Fumagalli, and F. Racimo. Detecting adaptive introgression in human evolution using convolutional neural networks. Elife, 10, May 2021.

Ryan N. Gutenkunst, Ryan D. Hernandez, Scott H. Williamson, and Carlos D. Bustamante. Inferring the joint demographic history of multiple populations from multidimensional SNP frequency data. PLOS Genetics, 5(10):1–11, 10 2009. doi: 10.1371/journal.pgen.1000695. URL https://doi.org/10.1371/journal.pgen.1000695.

Benjamin C. Haller and Philipp W. Messer. SLiM 3: Forward genetic simulations beyond the Wright-Fisher model. Molecular Biology and Evolution, 36(3):632–637, 2019.

Chad Harland, Carole Charlier, Latifa Karim, Nadine Cambisano, Manon Deckers, Myriam Mni, Erik Mullaart, Wouter Coppieters, and Michel Georges. Frequency of mosaicism points towards mutation-prone early cleavage cell divisions in cattle. bioRxiv, 2017. doi: 10.1101/079863. URL https://www.biorxiv.org/content/early/2017/06/29/079863.

K. Harris and R. Nielsen. Inferring demographic history from a spectrum of shared haplotype lengths. PLoS Genet, 9(6):e1003521, Jun 2013.

Kelley Harris and Rasmus Nielsen. The genetic cost of Neanderthal introgression. Genetics, 203(2):881–891, 06 2016. ISSN 1943-2631. doi: 10.1534/genetics.116.186890. URL https://doi.org/10.1534/genetics.116.186890.

M Hartfield, N Aagaard Poulsen, B Guldbrandtsen, and T Bataillon. Using singleton densities to detect recent selection in *Bos taurus*. Evolution Letters, 2022. doi: 10.1002/evl3.263. URL https://doi.org/10.1002/evl3.263.

Michael P Heaton, Timothy P L Smith, Derek M Bickhart, Brian L Vander Ley, Larry A Kuehn, Jonas Oppenheimer, Wade R Shafer, Fred T Schuetze, Brad Stroud, Jennifer C McClure, Jennifer P Barfield, Harvey D Blackburn, Theodore S Kalbfleisch, Kimberly M Davenport, Kristen L Kuhn, Richard E Green, Beth Shapiro, and Benjamin D Rosen. A reference genome assembly of Simmental cattle, *Bos taurus taurus*. Journal of Heredity, 112(2):184–191, 01 2021. ISSN 0022-1503. doi: 10.1093/jhered/esab002. URL https://doi.org/10.1093/jhered/esab002.

A. S. Hinrichs, D. Karolchik, R. Baertsch, G. P. Barber, G. Bejerano, H. Clawson, M. Diekhans, T. S. Furey, R. A. Harte, F. Hsu, J. Hillman-Jackson, R. M. Kuhn, J. S. Pedersen, A. Pohl, B. J. Raney, K. R. Rosenbloom, A. Siepel, K. E. Smith, C. W. Sugnet, A. Sultan-Qurraie, D. J. Thomas, H. Trumbower, R. J. Weber, M. Weirauch, A. S. Zweig, D. Haussler, and W. J. Kent. The UCSC Genome Browser Database: update 2006. Nucleic Acids Res, 34(Database issue):D590–598, Jan 2006.

Kevin L Howe, Premanand Achuthan, James Allen, Jamie Allen, Jorge Alvarez-Jarreta, M Ridwan Amode, Irina M Armean, Andrey G Azov, Ruth Bennett, Jyothish Bhai, Konstantinos Billis, Sanjay Boddu, Mehrnaz Charkhchi, Carla Cummins, Luca Da Rin Fioretto, Claire Davidson, Kamalkumar Dodiya, Bilal El Houdaigui, Reham Fatima, Astrid Gall, Carlos Garcia Giron, Tiago Grego, Cristina Guijarro-Clarke, Leanne Haggerty, Anmol Hemrom, Thibaut Hourlier, Osagie G Izuogu, Thomas Juettemann, Vinay Kaikala, Mike Kay, Ilias Lavidas, Tuan Le, Diana Lemos, Jose Gonzalez Martinez, José Carlos Marugán, Thomas Maurel, Aoife C McMahon, Shamika Mohanan, Benjamin Moore, Matthieu Muffato, Denye N Oheh, Dimitrios Paraschas, Anne Parker, Andrew Parton, Irina Prosovetskaia, Manoj P Sakthivel, Ahamed I Abdul Salam, Bianca M Schmitt, Helen Schuilenburg, Dan Sheppard, Emily Steed, Michal Szpak, Marek Szuba, Kieron Taylor, Anja Thormann, Glen Threadgold, Brandon Walts, Andrea Winterbottom, Marc Chakiachvili, Ameya Chaubal, Nishadi De Silva, Bethany Flint, Adam Frankish, Sarah E Hunt, Garth R IIsley, Nick Langridge, Jane E Loveland, Fergal J Martin, Jonathan M Mudge, Joanella Morales, Emily Perry, Magali Ruffier, John Tate, David Thybert, Stephen J Trevanion, Fiona Cunningham, Andrew D Yates, Daniel R Zerbino, and Paul Flicek. Ensembl 2021. Nucleic Acids Research, 49(D1):D884–D891, 11 2020. ISSN 0305-1048. doi: 10.1093/nar/gkaa942. URL https://doi.org/10.1093/nar/gkaa942.

PingHsun Hsieh, Krishna R Veeramah, Joseph Lachance, Sarah A Tishkoff, Jeffrey D Wall, Michael F Hammer, and Ryan N Gutenkunst. Whole genome sequence analyses of Western Central African Pygmy hunter-gatherers reveal a complex demographic history and identify candidate genes under positive natural selection. Genome Res., 26:279—-290, 2016.

PingHsun Hsieh, Vy Dang, Mitchell R Vollger, Yafei Mao, Tzu-Hsueh Huang, Philip C Dishuck, Carl Baker, Stuart Cantsilieris, Alexandra P Lewis, Katherine M Munson, et al. Evidence for opposing selective forces operating on human-specific duplicated tcaf genes in neanderthals and humans. Nature Communications, 12(1):5118, 2021.

Christian D. Huber, Arun Durvasula, Angela M. Hancock, and Kirk E. Lohmueller. Gene expression drives the evolution of dominance. Nat. Commun., 9(1):2750, 2018. ISSN 20411723. doi: 10.1038/s41467-018-05281-7. URL http://dx.doi.org/10.1038/s41467-018-05281-7.

T. H. Jukes and C. R. Cantor. Evolution of protein molecules. In H.N. Munro, editor, Mammalian Protein Metabolism, pages 21–132. Academic Press, New York, 1969.

P. D. Keightley, U. Trivedi, M. Thomson, F. Oliver, S. Kumar, and M. L. Blaxter. Analysis of the genome sequences of three *Drosophila melanogaster* spontaneous mutation accumulation lines. Genome Res, 19 (7):1195–1201, Jul 2009.

Jerome Kelleher, Alison M Etheridge, and Gilean McVean. Efficient coalescent simulation and genealogical analysis for large sample sizes. PLoS computational biology, 12(5):e1004842, 2016.

Katharine L. Korunes and Mohamed A. F. Noor. Gene conversion and linkage: effects on genome evolution and speciation. Molecular Ecology, 26(1):351–364, 2017. doi: https://doi.org/10.1111/mec.13736.

S. Kumar, M. Suleski, J. M. Craig, A. E. Kasprowicz, M. Sanderford, M. Li, G. Stecher, and S. B. Hedges. TimeTree 5: An Expanded Resource for Species Divergence Times. Mol Biol Evol, Aug 2022.

Christopher C. Kyriazis, Jacqueline A. Robinson, and Kirk E. Lohmueller. Using computational simulations to quantify genetic load and predict extinction risk. bioRxiv, 2022. doi: 10.1101/2022.08.12.503792. URL https://www.biorxiv.org/content/early/2022/08/15/2022.08.12.503792.

Greger Larson and Joachim Burger. A population genetics view of animal domestication. Trends in Genetics, 29(4):197–205, 2013.

Harris A. Lewin, Stephen Richards, Erez Lieberman Aiden, Miguel L. Allende, John M. Archibald, Miklós Bálint, Katharine B. Barker, Bridget Baumgartner, Katherine Belov, Giorgio Bertorelle, Mark L. Blaxter, Jing Cai, Nicolette D. Caperello, Keith Carlson, Juan Carlos Castilla-Rubio, Shu-Miaw Chaw, Lei Chen, Anna K. Childers, Jonathan A. Coddington, Dalia A. Conde, Montserrat Corominas, Keith A. Crandall, Andrew J. Crawford, Federica DiPalma, Richard Durbin, ThankGod E. Ebenezer, Scott V. Edwards, Olivier Fedrigo, Paul Flicek, Giulio Formenti, Richard A. Gibbs, M. Thomas P. Gilbert, Melissa M. Goldstein, Jennifer Marshall Graves, Henry T. Greely, Igor V. Grigoriev, Kevin J. Hackett, Neil Hall, David Haussler, Kristofer M. Helgen, Carolyn J. Hogg, Sachiko Isobe, Kjetill Sigurd Jakobsen, Axel Janke, Erich D. Jarvis, Warren E. Johnson, Steven J. M. Jones, Elinor K. Karlsson, Paul J. Kersey, Jin-Hyoung Kim, W. John Kress, Shigehiro Kuraku, Mara K. N. Lawniczak, James H. Leebens-Mack, Xueyan Li, Kerstin Lindblad-Toh, Xin Liu, Jose V. Lopez, Tomas Marques-Bonet, Sophie Mazard, Jonna A. K. Mazet, Camila J. Mazzoni, Eugene W. Myers, Rachel J. O’Neill, Sadye Paez, Hyun Park, Gene E. Robinson, Cristina Roquet, Oliver A. Ryder, Jamal S. M. Sabir, H. Bradley Shaffer, Timothy M. Shank, Jacob S. Sherkow, Pamela S. Soltis, Boping Tang, Leho Tedersoo, Marcela Uliano-Silva, Kun Wang, Xiaofeng Wei, Regina Wetzer, Julia L. Wilson, Xun Xu, Huanming Yang, Anne D. Yoder, and Guojie Zhang. The Earth BioGenome Project 2020: Starting the clock. Proceedings of the National Academy of Sciences, 119(4):e2115635118, 2022. doi: 10.1073/pnas.2115635118. URL https://www.pnas.org/doi/abs/10.1073/pnas.2115635118.

H. Li and R. Durbin. Inference of human population history from individual whole-genome sequences. Nature, 475(7357):493–496, Jul 2011.

X. Liu and Y. X. Fu. Corrigendum: Exploring population size changes using SNP frequency spectra. Nat Genet, 47(9):1099, Sep 2015.

Li Ma, Jeffrey R. O’Connell, Paul M. VanRaden, Botong Shen, Abinash Padhi, Chuanyu Sun, Derek M. Bickhart, John B. Cole, Daniel J. Null, George E. Liu, Yang Da, and George R. Wiggans. Cattle sex-specific recombination and genetic control from a large pedigree analysis. PLOS Genetics, 11(11):1–24, 11 2015. doi: 10.1371/journal.pgen.1005387. URL https://doi.org/10.1371/journal.pgen.1005387.

Xin Ma, Joanna L. Kelley, Kirsten Eilertson, Shaila Musharoff, Jeremiah D. Degenhardt, André L. Martins, Tomas Vinar, Carolin Kosiol, Adam Siepel, Ryan N. Gutenkunst, and Carlos D. Bustamante. Population genomic analysis reveals a rich speciation and demographic history of orang-utans (*Pongo pygmaeus* and *Pongo abelii*). PLoS One, 8(10):e77175, oct 2013. ISSN 1932-6203. doi: 10.1371/journal.pone.0077175. URL http://dx.plos.org/10.1371/journal.pone.0077175.

I M MacLeod, D M Larkin, H A Lewin, B J Hayes, and M E Goddard. Inferring demography from runs of homozygosity in whole-genome sequence, with correction for sequence errors. Molecular Biology and Evolution, 30(9):2209–2223, 07 2013. ISSN 0737-4038. doi: 10.1093/molbev/mst125. URL https://doi.org/10.1093/molbev/mst125.

I M MacLeod, B J Hayes, and M E Goddard. The Effects of Demography and Long-Term Selection on the Accuracy of Genomic Prediction with Sequence Data. Genetics, 198(4):1671–1684, 09 2014. ISSN 1943-2631. doi: 10.1534/genetics.114.168344. URL https://doi.org/10.1534/genetics.114.168344.

B O Makanjuola, F Miglior, E A Abdalla, C Maltecca, F S Schenkel, and C F Baes. Effect of genomic selection on rate of inbreeding and coancestry and effective population size of Holstein and Jersey cattle populations. Journal of Dairy Science, 2020. doi: 10.3168/jds.2019-18013. URL https://doi.org/10.3168/jds.2019-18013.

G. McVicker, D. Gordon, C. Davis, and P. Green. Widespread genomic signatures of natural selection in hominid evolution. PLoS Genet, 5(5):e1000471, May 2009.

T H E Meuwissen, B J Hayes, and M E Goddard. Prediction of total genetic value using genome-wide dense marker maps. Genetics, 157(4):1819–1829, 04 2001. ISSN 1943-2631. doi: 10.1093/genetics/157.4.1819. URL https://doi.org/10.1093/genetics/157.4.1819.

A. Miles, N. J. Harding, G. Botta, C. S. Clarkson, T. Antao, K. Kozak, D. R. Schrider, A. D. Kern, S. Redmond, I. Sharakhov, R. D. Pearson, C. Bergey, M. C. Fontaine, M. J. Donnelly, M. K. N. Lawniczak, D. P. Kwiatkowski, M. J. Donnelly, D. Ayala, N. J. Besansky, A. Burt, B. Caputo, A. Della Torre, M. C. Fontaine, H. C. J. Godfray, M. W. Hahn, A. D. Kern, D. P. Kwiatkowski, M. K. N. Lawniczak, J. Midega, D. E. Neafsey, S. O’Loughlin, J. Pinto, M. M. Riehle, I. Sharakhov, K. D. Vernick, D. Weetman, C. S. Wilding, B. J. White, A. D. Troco, J. Pinto, A. Diabaté, S. O’Loughlin, A. Burt, C. Costantini, K. R. Rohatgi, N. J. Besansky, N. Elissa, J. Pinto, B. Coulibaly, M. M. Riehle, K. D. Vernick, J. Pinto, J. Dinis, J. Midega, C. Mbogo, P. Bejon, C. S. Wilding, D. Weetman, H. D. Mawejje, M. J. Donnelly, D. Weetman, C. S. Wilding, M. J. Donnelly, J. Stalker, K. Rockett, E. Drury, D. Mead, A. Jeffreys, C. Hubbart, K. Rowlands, A. T. Isaacs, D. Jyothi, C. Malangone, P. Vauterin, B. Jeffery, I. Wright, L. Hart, K. Kluczy?ski, V. Cornelius, B. MacInnis, C. Henrichs, R. Giacomantonio, D. P. Kwiatkowski, V. Cornelius, B. MacInnis, C. Henrichs, R. Giacomantonio, and D. P. Kwiatkowski. Genetic diversity of the African malaria vector Anopheles gambiae. Nature, 552(7683):96–100, 12 2017.

Valeria Montano. Coalescent inferences in conservation genetics: Should the exception become the rule? Biol. Lett., 12(6), 2016. ISSN 1744957X. doi: 10.1098/rsbl.2016.0211.

Michael W. Nachman. Variation in recombination rate across the genome: Evidence and implications. Curr. Opin. Genet. Dev., 12(6):657–663, 2002. ISSN 0959437X. doi: 10.1016/S0959-437X(02)00358-1.

Miguel Navascués and Brent C Emerson. Elevated substitution rate estimates from ancient DNA: model violation and bias of Bayesian methods. Molecular Ecology, 18(21):4390–4397, 2009.

Dominic Nelson, Jerome Kelleher, Aaron P. Ragsdale, Claudia Moreau, Gil McVean, and Simon Gravel. Accounting for long-range correlations in genome-wide simulations of large cohorts. PLOS Genetics, 16 (5):1–12, 05 2020. doi: 10.1371/journal.pgen.1008619. URL https://doi.org/10.1371/journal.pgen.1008619.

J Obšteter, J Jenko, and G Gorjanc. Genomic selection for any dairy breeding program via optimized investment in phenotyping and genotyping. Frontiers in Genetics, 12, 2021. doi: 10.3389/fgene.2021. 637017. URL https://www.frontiersin.org/article/10.3389/fgene.2021.637017.

March Pombi, Aram D. Stump, Allesandra Della Torre, and Nora J. Besansky. Variation in recombination rate across the X chromosome of *Anopheles gambiae*. The American Journal of Tropical Medicine and Hygiene., 75(5):901–903, 2006. doi: https://doi.org/10.4269/ajtmh.2006.75.901. URL https://www.ajtmh.org/view/journals/tpmd/75/5/article-p901.xml.

Aaron P. Ragsdale, Dominic Nelson, Simon Gravel, and Jerome Kelleher. Lessons learned from bugs in models of human history. The American Journal of Human Genetics, 107(4):583–588, 2020. ISSN 0002-9297. doi: https://doi.org/10.1016/j.ajhg.2020.08.017. URL https://www.sciencedirect.com/science/article/pii/S000292972030286X.

Arang Rhie, Shane A. McCarthy, Olivier Fedrigo, Joana Damas, Giulio Formenti, Sergey Koren, Marcela Uliano-Silva, William Chow, Arkarachai Fungtammasan, Juwan Kim, Chul Lee, Byung June Ko, Mark Chaisson, Gregory L. Gedman, Lindsey J. Cantin, Francoise Thibaud-Nissen, Leanne Haggerty, Iliana Bista, Michelle Smith, Bettina Haase, Jacquelyn Mountcastle, Sylke Winkler, Sadye Paez, Jason Howard, Sonja C. Vernes, Tanya M. Lama, Frank Grutzner, Wesley C. Warren, Christopher N. Balakrishnan, Dave Burt, Julia M. George, Matthew T. Biegler, David Iorns, Andrew Digby, Daryl Eason, Bruce Robertson, Taylor Edwards, Mark Wilkinson, George Turner, Axel Meyer, Andreas F. Kautt, Paolo Franchini, H. William Detrich III, Hannes Svardal, Maximilian Wagner, Gavin J. P. Naylor, Martin Pippel, Milan Malinsky, Mark Mooney, Maria Simbirsky, Brett T. Hannigan, Trevor Pesout, Marlys Houck, Ann Misuraca, Sarah B. Kingan, Richard Hall, Zev Kronenberg, Ivan Sović, Christopher Dunn, Zemin Ning, Alex Hastie, Joyce Lee, Siddarth Selvaraj, Richard E. Green, Nicholas H. Putnam, Ivo Gut, Jay Ghurye, Erik Garrison, Ying Sims, Joanna Collins, Sarah Pelan, James Torrance, Alan Tracey, Jonathan Wood, Robel E. Dagnew, Dengfeng Guan, Sarah E. London, David F. Clayton, Claudio V. Mello, Samantha R. Friedrich, Peter V. Lovell, Ekaterina Osipova, Farooq O. Al-Ajli, Simona Secomandi, Heebal Kim, Constantina The-ofanopoulou, Michael Hiller, Yang Zhou, Robert S. Harris, Kateryna D. Makova, Paul Medvedev, Jinna Hoffman, Patrick Masterson, Karen Clark, Fergal Martin, Kevin Howe, Brian P. Flicek, Paul Walenz, Woori Kwak, Hiram Clawson, Mark Diekhans, Luis Nassar, Benedict Paten, Robert H. S. Kraus, Andrew J. Crawford, M. Thomas P. Gilbert, Guojie Zhang, Byrappa Venkatesh, Robert W. Murphy, Klaus-Peter Koepfli, Beth Shapiro, Warren E. Johnson, Federica Di Palma, Tomas Marques-Bonet, Emma C. Teeling, Tandy Warnow, Jennifer Marshall Graves, Oliver A. Ryder, David Haussler, Stephen J. O’Brien, Jonas Korlach, Harris A. Lewin, Kerstin Howe, Eugene W. Myers, Richard Durbin, Adam M. Phillippy, and Erich D. Jarvis. Towards complete and error-free genome assemblies of all vertebrate species. Nature, 592(7856):737–746, 2021. ISSN 1476-4687. doi: 10.1038/s41586-021-03451-0.

J. Robinson, C. C. Kyriazis, S. C. Yuan, and K. E. Lohmueller. Deleterious Variation in Natural Populations and Implications for Conservation Genetics. Annu Rev Anim Biosci, 11:93–114, Feb 2023.

Benjamin D Rosen, Derek M Bickhart, Robert D Schnabel, Sergey Koren, Christine G Elsik, Elizabeth Tseng, Troy N Rowan, Wai Y Low, Aleksey Zimin, Christine Couldrey, Richard Hall, Wenli Li, Arang Rhie, Jay Ghurye, Stephanie D McKay, Françoise Thibaud-Nissen, Jinna Hoffman, Brenda M Murdoch, Warren M Snelling, Tara G McDaneld, John A Hammond, John C Schwartz, Wilson Nandolo, Darren E Hagen, Christian Dreischer, Sebastian J Schultheiss, Steven G Schroeder, Adam M Phillippy, John B Cole, Curtis P Van Tassell, George Liu, Timothy P L Smith, and Juan F Medrano. De novo assembly of the cattle reference genome with single-molecule sequencing. GigaScience, 9(3), 03 2020. ISSN 2047-217X. doi: 10.1093/gigascience/giaa021. URL https://doi.org/10.1093/gigascience/giaa021.giaa021.

Stephan Schiffels and Ke Wang. MSMC and MSMC2: The Multiple Sequentially Markovian Coalescent, pages 147–166. Springer US, New York, NY, 2020. ISBN 978-1-0716-0199-0. doi: 10.1007/978-1-0716-0199-0_7. URL https://doi.org/10.1007/978-1-0716-0199-0_7.

Daniel R Schrider. Background selection does not mimic the patterns of genetic diversity produced by selective sweeps. Genetics, 216(2):499–519, 2020.

Daniel R. Schrider and Andrew D. Kern. Supervised machine learning for population genetics: A new paradigm. Trends Genet., 34(4):301–312, 2018. ISSN 13624555. doi: 10.1016/j.tig.2017.12.005. URL http://dx.doi.org/10.1016/j.tig.2017.12.005.

M. V. Sharakhova, M. P. Hammond, N. F. Lobo, J. Krzywinski, M. F. Unger, M. E. Hillenmeyer, R. V. Bruggner, E. Birney, and F. H. Collins. Update of the *Anopheles gambiae* PEST genome assembly. Genome Biol, 8(1):R5, 2007.

Fran Supek and Ben Lehner. Scales and mechanisms of somatic mutation rate variation across the human genome. DNA Repair, 81:102647, 2019. ISSN 1568-7864. doi: https://doi.org/10.1016/j.dnarep.2019.102647. URL https://www.sciencedirect.com/science/article/pii/S1568786419302009. Cuttingedge Perspectives in Genomic Maintenance VI.

A Talenti, J Powell, J D Hemmink, E A J Cook, D Wragg, S Jayaraman, E Paxton, C Ezeasor, E T Obishakin, E R Agusi, A Tijjani, K Marshall, A Fisch, B R Ferreira, A Qasim, U Chaudhry, P Wiener, P Toye, L J Morrison, T Connelley, and J G D Prendergast. A cattle graph genome incorporating global breed diversity. Nature Communications, 2022. doi: 10.1038/s41467-022-28605-0. URL https://doi.org/10.1038/s41467-022-28605-0.

João C. Teixeira and Christian D. Huber. The inflated significance of neutral genetic diversity in conservation genetics. Proc. Natl. Acad. Sci. U. S. A., 118(10):1–10, 2021. ISSN 10916490. doi: 10.1073/pnas.2015096118.

J. A. Tennessen, A. W. Bigham, T. D. O’Connor, W. Fu, E. E. Kenny, S. Gravel, S. McGee, R. Do, X. Liu, G. Jun, H. M. Kang, D. Jordan, S. M. Leal, S. Gabriel, M. J. Rieder, G. Abecasis, D. Altshuler, D. A. Nickerson, E. Boerwinkle, S. Sunyaev, C. D. Bustamante, M. J. Bamshad, and J. M. Akey. Evolution and functional impact of rare coding variation from deep sequencing of human exomes. Science, 337(6090): 64–69, Jul 2012.

Kosuke M. Teshima, Graham Coop, and Molly Przeworski. How reliable are empirical genomic scans for selective sweeps? Genome Res., 16(6):702–712, 2006. ISSN 10889051. doi: 10.1101/gr.5105206.

C. M. Thomas and K. M. Nielsen. Mechanisms of, and barriers to, horizontal gene transfer between bacteria. Nat Rev Microbiol, 3(9):711–721, Sep 2005.

P M VanRaden. Symposium review: How to implement genomic selection. Journal of Dairy Science, 103(6):5291–5301, 2020. ISSN 0022-0302. doi: https://doi.org/10.3168/jds.2019-17684. URL https://www.sciencedirect.com/science/article/pii/S002203022030309X.

S. Wielgoss, J. E. Barrick, O. Tenaillon, S. Cruveiller, B. Chane-Woon-Ming, C. Medigue, R. E. Lenski, and D. Schneider. Mutation rate inferred from synonymous substitutions in a long-term evolution experiment with *Escherichia coli*. G3 (Bethesda), 1(3):183–186, Aug 2011.

E. Wijnker, G. Velikkakam James, J. Ding, F. Becker, J. R. Klasen, V. Rawat, B. A. Rowan, D. F. de Jong, C. B. de Snoo, L. Zapata, B. Huettel, H. de Jong, S. Ossowski, D. Weigel, M. Koornneef, J. J. Keurentjes, and K. Schneeberger. The genomic landscape of meiotic crossovers and gene conversions in *Arabidopsis thaliana*. Elife, 2:e01426, Dec 2013.

Liangbiao Zheng, Mark Q Benedict, Anton J Cornel, Frank H Collins, and Fotis C Kafatos. An integrated genetic map of the African human malaria vector mosquito, Anopheles gambiae. Genetics, 143(2):941–952, 1996.

Ying Zhou, Xiaowen Tian, Brian L Browning, and Sharon R Browning. POPdemog: visualizing population demographic history from simulation scripts. Bioinformatics, 34(16):2854–2855, 03 2018. ISSN 1367-4803. doi: 10.1093/bioinformatics/bty184. URL https://doi.org/10.1093/bioinformatics/bty184.

